# Replication rate-information storage trade-off shapes genome architecture across domains

**DOI:** 10.1101/2025.08.07.669222

**Authors:** Parthasarathi Sahu, Sashikanta Barik, Koushik Ghosh, Hemachander Subramanian

## Abstract

Genome size varies widely among organisms, from compact genomes of bacteria to vast and complex genomes of eukaryotes. In this study, we theoretically identify the evolutionary pressures that may have driven this divergence in genome size. We use a parameter-free model to study genome size evolution under selection pressure to minimize replication time and maximize information storage capacity. We substantiate this choice by demonstrating a correlation between replication time and genome size using literature data. We show that bacteria tend to reduce genome size, constrained by a single replication origin, while eukaryotes expand their genomes by incorporating multiple replication origins. We propose a connection between genome size and cellular energetics, suggesting that endosymbiotic organelles, mitochondria and chloroplasts, evolutionarily regulate the number of replication origins, thereby influencing genome size in eukaryotes. We substantiate this claim by showing a correlation between organelle count and genome size data. We argue that nucleotide skews, present in nearly all genomes studied, and are used to identify replication origins, directly influence DNA unzipping kinetics, and hence its replication time. This connection enables us to derive nearly universal observations, such as Chargaff’s second parity rule and replichore symmetrization, as adaptive consequences. We argue that high skews lead to faster replication, and substantiate it by showing a correlation between skews and replication speed in bacteria. The model reproduces more evolutionary genomic observations, such as a general preference for deletions over insertions across domains, and elongation and high variance of genome size under reduced selection pressure for replication rate, an integral component of C-value paradox. We highlight the possibility of regulation of the firing of latent replication origins in response to cues from the extracellular environment, leading to the regulation of cell cycle rates in multicellular eukaryotes.

**Significance Statement:** Understanding the forces shaping genome architecture is a long-standing challenge in evolutionary biology. Our study demonstrates that the balance between replication speed, influenced by nucleotide skews, and information storage, constrained by cellular energetics, drives the divergence in genome size between bacteria and eukaryotes. By quantifying selection pressure as the ratio of replication time to genomic information storage capacity, we show that this pressure enforces adaptive constraints, giving rise to observed features such as symmetric replichores and Chargaff’s second parity rule. These insights not only help us resolve an enduring evolutionary puzzle, but also offer a unified framework linking genome organization, cellular specialization, and even potential mechanisms underlying carcinogenesis.

## Introduction

Life on Earth began approximately 3.7 billion years ago and evolved from simpler forms to complex and diverse organisms observed today, shaped by various selection pressures [1, 2]. Organisms are broadly classified into three groups: bacteria, archaea, and eukaryotes. Bacteria and archaea are characterized by simpler intracellular structures, whereas eukaryotes are generally more complex, with defining characteristics such as endosymbiotic relationships, nuclear membranes, huge variance in genome size, etc. [3, 4, 5]. Despite emerging earlier in Earth’s history, bacteria and archaea maintain smaller genomes and show less intracellular structural complexity compared to eukaryotes [6, 7, 8]. The constraints limiting the evolution of such complexity in bacteria and archaea are debated [9, 10, 11].

The tendency of bacteria and archaea to maintain compact genomes is extensively modeled, with models constructed to include impacts of population size, environmental perturbations, and selection for metabolic efficiency under nutrient limitation [6, 12, 13, 14]. The evolutionary forces shaping eukaryotic genome size remain comparatively under-explored. In eukaryotes, current frameworks have focused on the impact of mutational mechanisms [15, 16] and energetic constraints on genome size [17].

One of the major differences between these two groups is genome size, with bacteria and archaea having a 20fold variation in genome size, whereas eukaryotic genomes vary 200,000-fold [18, 19]. It is also evident that this dramatic spread in eukaryotic genome size doesn’t correspond to organismal complexity [20]. The lack of correlation of genome size with organismal complexity is termed as ‘C-value paradox’ [21]. There are two classes of explanation for this paradox: (a) mutational pressure theories [22, 23] and (b) optimal DNA theories [24, 25]. Under the first class, junk and selfish DNA are considered responsible for the expansion of the genome size in eukaryotes. It is argued that the inevitable expansion of efficient self-replicators would produce a constant pressure to increase genome size, and this expansion is curtailed only when the additional genetic material becomes excessively burdensome for the host cell [26]. In the second class, phenotypic constraints, such as cell volume, evolutionarily select a specific genome size.

Despite extensive studies, the reason for such a dramatic divergence of genome size among the three domains of life is still debated. In this study, we use a very simple, barebones, parameter-free model that incorporates the influence of two primary evolutionary forces: *faster replication and enhanced information storage capacity*, to study the evolution of genome size across these three domains of life. We show that the genome sizes of bacteria, archaea, and eukaryotes diverge under the same selection pressure, if we restrict bacteria to have a single replication origin, archaea to have a few, while allowing eukaryotes to have multiple origins. We argue that this differentiation stems from access to the energy supply of mitochondria (or chloroplasts), as evidenced by the correlation we observe between the number of mitochondria (or chloroplasts) and the size of the genome in eukaryotes. Surprisingly, the model also reproduces multiple other observations that hold for nearly all free-living species, such as the equality of purines and pyrimidines on a single strand of DNA, called Chargaff’s second parity rule (PR-2), replichore length symmetrization (see below), a preference for deletions over insertion mutations, and a huge variance in the genome size seen in eukaryotes, a component of the C-Value paradox. These non-trivial outcomes arise primarily from *our inclusion of the influence of nucleotide skews on DNA unzipping kinetics*, which the model uses to detect and preserve replication origins during evolution, an approach not used in genome evolution models.

Genome size evolution is often portrayed as primarily neutral, driven by selfish elements proliferating under weak selection. However, a substantial and growing body of literature supports an adaptive interpretation of genome expansion, [27, 28, 29, 30, 31, 32, 33, 34, 35], as reviewed in literature [36, 33, 24]. Notably, the role of so-called “junk DNA” remains incompletely resolved, and its functional and informational significance is far from fully known [37, 38, 39, 40, 41, 42, 43]. In this work, we treat genome size, which includes coding and non-coding parts, as an adaptive trait, where increases in genome size may confer benefits such as enhanced regulatory potential, developmental flexibility, and environmental responsiveness, and we use *information storage capacity* as a catch-all term to include these adaptive gains.

### The Model

To study the effect of the aforementioned selection pressures on genome length, i.e., faster replication and higher information storage capacity, we utilize a simple model where a pool of *N* identical sequences is evolved over *m* generations under these selection pressures. The initial sequence pool consists of *N* identical sequences, containing purines or pyrimidines, homogeneously across the sequence, with all purines on one strand and all pyrimidines on the complementary strand (e.g., 5′-RRRRRRR-3′*/*3′-YYYYYYY-5′). We will later see that this specific, unnatural choice of initial pool is to demonstrate the effect of selection pressure on this sub-optimal choice, leading to the evolution of genomic architectures closer to those observed in nature. Each generation involves two major steps: (i) replication of *N* sequences and mutation of all daughter sequences in the pool, and (ii) selection of half of the sequences in the pool based on their ability to satisfy the above two selection pressures. These two steps are repeated *m* times, and the time-evolution of the sequence length, averaged over all the *N* sequences, is recorded for every generation, for subsequent analysis.

### Mutation

We implement large-scale genomic mutations through random deletions or duplications of regions in the daughter sequence that comprise 5% to 10% of the total genome length (Fig. 2). These large-scale mutations mimic transpositions and represent well-documented drivers of evolutionary processes [6, 44, 45, 46, 11]. The size and location of these mutations are chosen stochastically. During a mutation involving duplication, the duplicating fragment is randomly chosen from either the original strand or the complementary strand. This mechanism ensures that over generations, each strand can have varying amounts of both purines and pyrimidines, even though the initial pool sequences are composed entirely of either purines or pyrimidines. In each generation, every sequence is replicated, and the replicated sequence in the pool undergoes a single mutation, i.e., a duplication or deletion. Following this, the sequence pool is expanded to include *N* replicated and mutated sequences along with the *N* unmutated sequences of the previous generation, resulting in a total of 2*N* sequences.

**Figure 1.**
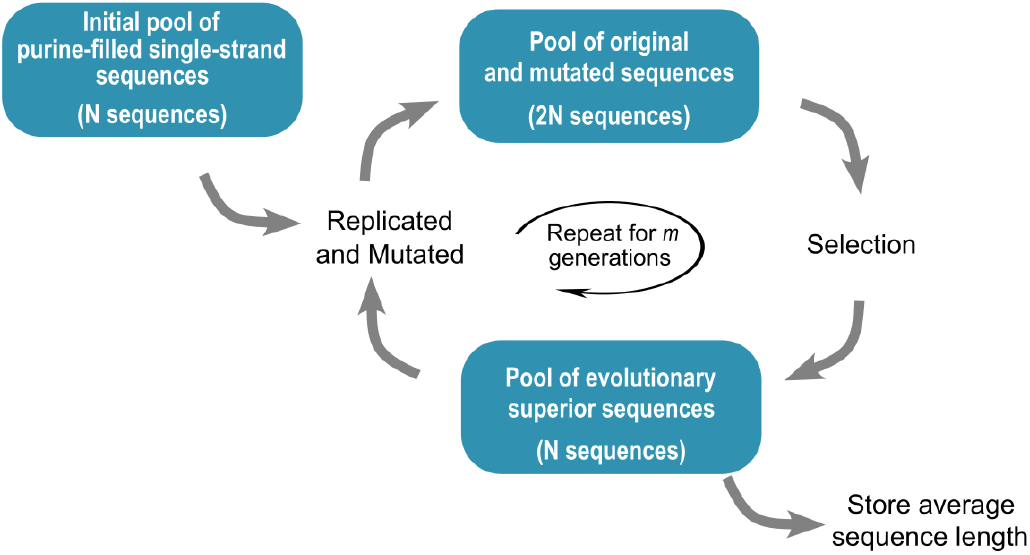
Algorithm of the model. Evaluation of the impact of the two selection pressures, fast replication and high information storage capacity, on the genome length of an organism. An initial pool of *N* sequences is evolved over *m* generations that involve two recurring steps: replication and mutation of all the sequences in the pool, and applying selection pressure to extract the fittest sequences for the next generation. An initial pool of *N* identical sequences, composed of all purines or all pyrimidines, is replicated, and the daughter sequences are mutated, producing a pool of 2*N* sequences. Selection acts on this pool, removing *N* less-fit sequences that do not satisfy the selection pressure adequately. This replication-selection cycle is repeated *m* times, and the average genome length at every generation is recorded.

**Figure 2.**
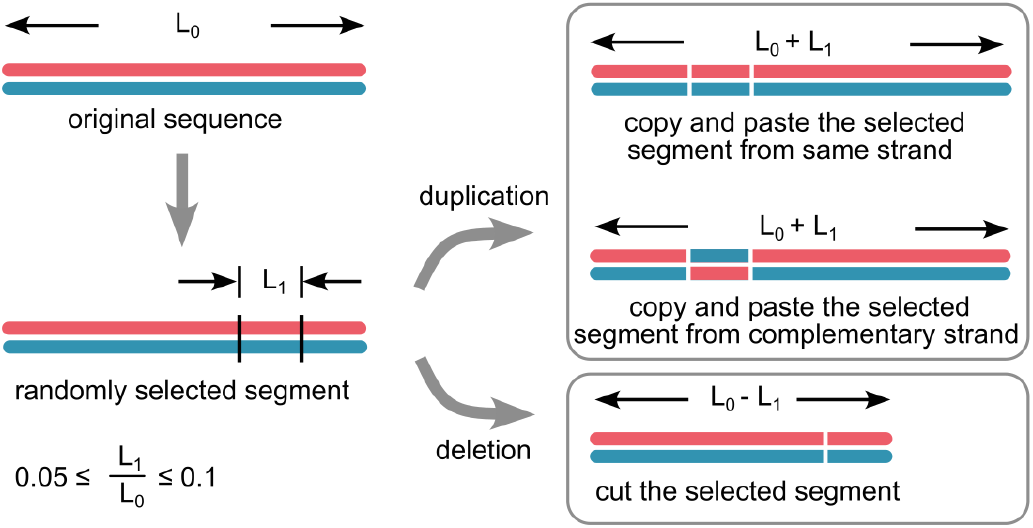
Mutation of a given genome. A DNA double strand, from the initial pool, with a homogeneous distribution of purines on one strand (red) and pyrimidines on its complement (blue). A mutation involves either a deletion or a duplication of a segment of a random length of 5-10% of the total genome length, chosen at a random location of the daughter genome. The mutation results in either a decrease or an increase in the length of the genome by 5-10%. The duplicated fragment can be from either strand, thereby altering the composition of purines/pyrimidines in each strand.

### Selection

Following mutations, *N* sequences are selected from the pool of 2*N* for the next generation based on their ability to satisfy the selection pressure. The selection pressure is quantified through a factor *γ*, defined as

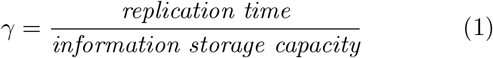

We selected these two evolutionary pressures to model two opposing evolutionary forces: an upward drive to increase genome length (information storage capacity) and a downward drive to minimize replication time, and study their trade-off. Information storage capacity, defined as total genome length, encompasses all adaptive gains from genome expansion. Genome growth often arises from non-coding regions, which typically precede the de novo emergence of functional genes, suggesting that genome expansion is a prerequisite for novel gene evolution [47, 48, 49, 50, 51, 52, 53, 54]. Replication time is chosen as another selection pressure on genome size, since it has been reported to inversely correlate with cellular growth rate [55, 56, 57, 58, 59, 60]. This is also supported by our analysis of 47 data points (with distinct ploidy) from 44 species [61](supplementary data), which reveals a significant correlation between DNA content and S-phase duration (Fig. 3), further validating replication time as a biologically relevant selection pressure.

**Figure 3.**
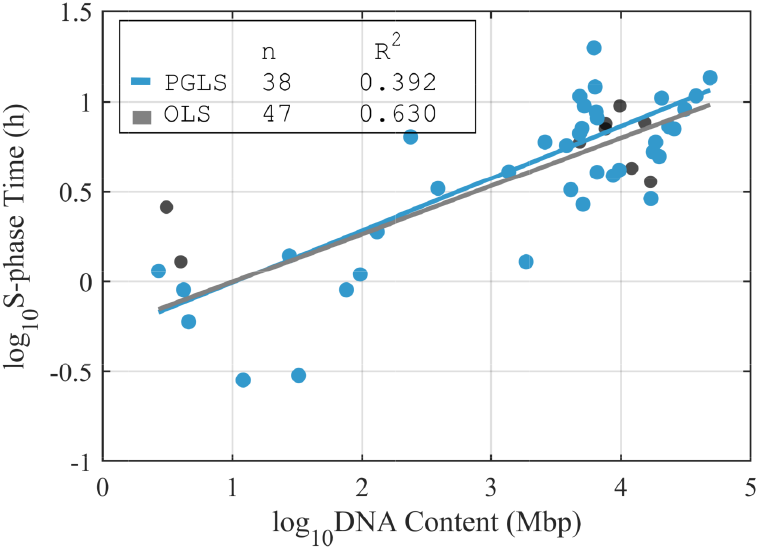
Correlation between genome size and replication time. The relationship between DNA content and S-phase duration in the cell cycle, in 47 data points across 44 organisms from diverse domains [61], suggests that replication dynamics may influence genome size evolution. This finding supports the inclusion of replication time as a selection pressure in our genome evolution model. Notably, while genome sizes span nearly 10^5^-fold, S-phase durations vary by only ~ 10^2^-fold, indicating that the organism with the largest genome in this dataset likely achieves at least a 10^3^-fold increase in replication parallelization across replichores compared with the organism having the smallest genome, assuming similar replication fork progression speeds. If the replication parallelization is complete, with all replichores replicating simultaneously, this correlation would disappear. The persistence of correlation between replication time and genome size implies incomplete parallelization and the imposition of a certain temporal order in the firing of replication origins as commonly observed.

To quantify information storage capacity, we utilize the metric of total genome length. Similarly, as a proxy for the replication time of the entire genome, we consider the length of the longest *replichore*, where replichores are defined as disjoint segments of the genome that replicate independently of each other. This substitution is based on the following considerations: (a) Replichores replicate simultaneously and independently, parallelizing the replication process [62, 63]. Although the firing of multiple replication origins across the genome is generally not synchronized [64, 65], we assume ideal conditions, where origins fire and replication progress across each replichore independently and simultaneously. (b) For simplicity, it is assumed here that the replication speed of a replichore is constant throughout the length of the replichore, although the speed depends on the sequence and the availability of activated nucleotides, in general [66]. This simplifying assumption enables us to use the replichore length as a replacement for the replication time of the whole genome, in our evaluation of *γ*. Therefore, with the above assumptions, we redefine equation (1) as

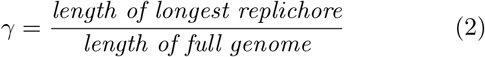

In the selection process, the factor *γ* is calculated for all the 2*N* (parent and mutated daughter) sequences in the pool, and the *N* sequences with the lowest values of *γ* are extracted as the fittest sequences and are carried forward to the next generation.

### Identification of replichores

To identify replichores in a genome, we first locate replication origins and termini by analyzing base composition asymmetry or nucleotide skews, i.e., the excess of G over C and A over T on any given stretch of a single strand, across the genome. This local violation of Chargaff’s second parity rule [67] is regularly utilized for *in-silico* prediction of replication origins and termini, by equating the locations of peaks and valleys in the skew plot to the replication origins and termini [63, 68, 69, 70, 71].

While this asymmetry in base composition has traditionally been attributed to mutational biases between the leading and lagging strands [68], we argue, based on our earlier model results [63], that the V-shaped purine-pyrimidine skews *cause* local strand melting, leading to the creation of replication origins. Our approach is motivated by previous DNA unzipping simulations, which demonstrated that an asymmetric nearest-neighbor kinetic influence of base pairs facilitates local double-strand melting at sites of compositional asymmetry, thereby initiating replication origins [63]. This kinetic phenomenon,termed *Asymmetric Cooperativity*, had been proposed to explain some key evolutionary features of DNA, including the advantage of unidirectional strand synthesis [72], antiparallel strand orientation [62], the necessity of a four-nucleotide genetic alphabet [73], and the requirement of compositional skew for local strand separation during origin activation [63]. Within the framework of the Asymmetric Cooperative Model, double-stranded DNA unzipping occurs more rapidly in regions with pronounced purine–pyrimidine (RY) skew, producing a characteristic “V-shaped” cumulative skew profile (Fig. 4(a)), and disruption of this structural asymmetry reduces unzipping efficiency (Figure 4(b)) [62, 63]. Experimental findings support this mechanism: Guilbaud et al. [74] reported that efficient replication origins exhibit strong GC skew, whereas dormant origins show weaker skew; Wanrooij et al. [75] further demonstrated that point mutations diminishing this V-shaped skew reduce origin activity, even when stable secondary structures (e.g., stem-loops) persist with an inverted “∧-shaped” skew profile.

**Figure 4.**
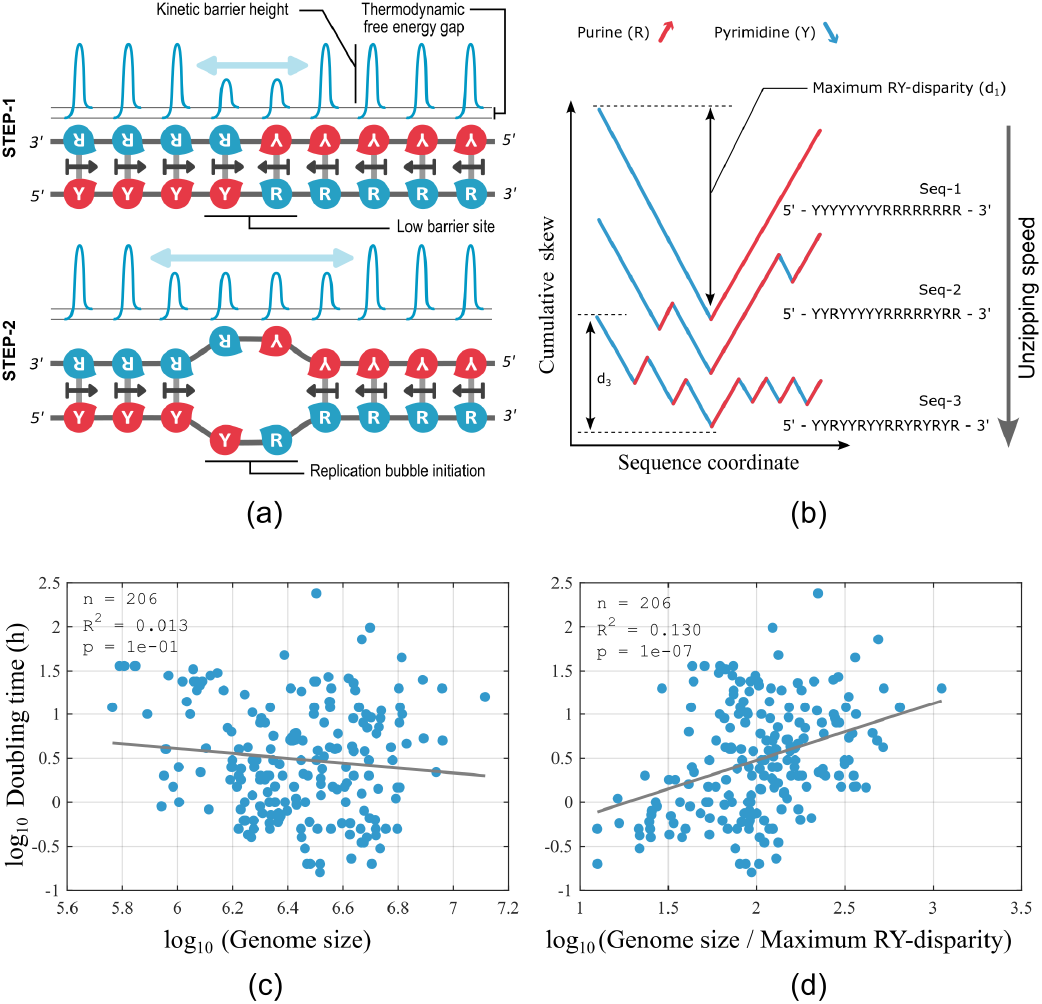
Purine–pyrimidine compositional asymmetry influences replication speed. (a) Kinetic barrier heights for base-pair hybridization/dehybridization are shown in blue above each base pair, illustrating the asymmetric cooperative effect proposed by the Asymmetric Cooperative Model. This mechanism explains local melting of double-stranded DNA at V-shaped RY-skew sites. (b) Skew profiles of three sequences with varying maximum local RY-disparity are shown. The Asymmetric Cooperativity model predicts that stronger skew promotes faster local DNA unzipping, leading to shorter replication times. (c) Without accounting for this sequence property, genome size shows no correlation with doubling time across 206 bacterial species under rapid growth conditions. (d) When genome sizes are normalized by their corresponding maximum local RY-skew, a mild but significant correlation emerges between doubling time and the logarithm of skew-normalized genome size, supporting the role of sequence structure (RY-skew) in replication speed.

Furthermore, in agreement with these findings, we quantified the *maximum local RY-disparity* of each genome by identifying the sequence window with the highest enrichment of purines over pyrimidines (or vice versa), defining this excess as the maximum local RY-disparity (Fig.4(b)). We then normalized genome size by this measure, referring to the resulting value as the skew-normalized genome size. Using this metric, analysis of 206 bacterial genomes revealed a mild yet significant (*p <* 10^−7^) correlation between the logarithm of skew-normalized genome size and the logarithm of cellular doubling time under rapid growth conditions (Fig.4(d)). While previous studies reported no association between bacterial doubling time and genome size [11, 76], our reanalysis of the same dataset [77] demonstrates that accounting for RY-skew uncovers this relationship. These findings suggest that purine–pyrimidine compositional asymmetry modulates replication kinetics. The sequence data used in the analysis were retrieved from the GenBank database (https://www.ncbi.nlm.nih.gov/genbank/), with accession numbers listed in Supplementary Table 1. The MATLAB code used to compute maximum local RY-disparity is provided in the Supplementary Material. Based on these findings, we adopt RY-skew switching (V-shaped local skew profile) as the primary criterion for replication origin identification in our model. However, the current implementation does not yet incorporate the direct effect of skew on replication speed, which will be added in future model extensions.

**Table 1.** Phylogenetic signal strength (*K*) from PGLS analysis for mitochondria and chloroplast datasets.

| Mitochondrial dataset |  | Chloroplast dataset |  |
| --- | --- | --- | --- |
| Trait | $K$ | Trait | $K$ |
| Mitochondria Count | 0.3741 | Chloroplast Count | 0.7217 |
| Genome Size | 0.5502 | Genome Size | 0.2738 |

In this study, we have used the purine-pyrimidine (RY) cumulative skew, *W*, of the sequences [78, 79], where *W*_*RY*_ (*n*) is defined as 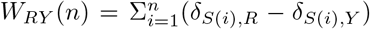. Here,*S* is a genomic sequence of length *L* bp, composed of four nucleotides, classified into two groups: *R* = {*G, A*}, *Y* = {*C, T*}, and *n* = 1 … *L*. We have taken 1000 purine-filled single strands of length 1024 bp each as the initial sequence pool. These sequences are allowed to mutate, by accrual or deletion of genome fragments, leading to local distortions in the skew in each generation. In order to avoid identifying small-scale skew variations (peaks and valleys smaller than a certain length scale) as origins or termini, since these are not identified as origins or termini by origin-finding algorithms [79, 80, 81, 82], we concentrate only on large-scale skew variations and ignore small-scale ones. We use wavelet transforms to filter out these origins and termini resulting from small-scale variations, by down-sampling the genome sequence of length *L* to a length of *L/*2^*ω*^, where *ω* is the wavelet level. To ensure that mutating fragments are not smaller than the wavelet compression scale *ω* and thus go unnoticed by the selection pressure, we choose a *ω* such that the smallest mutating fragment is larger than the compression factor; i.e., we impose the condition 5% *of L >* 2^*ω*^, where *L* is the full genome length. It should be emphasized that the qualitative divergence of bacterial and eukaryotic genome lengths does not depend on the wavelet level used in this down-sampling procedure (Supplementary Data 2). Once the origins and termini of the replication are identified, the lengths of the replichores were measured as the distances between neighboring origins and termini, the largest of which is chosen for the calculation of *γ*. A detailed description of the methodology for identifying replication origins and measuring replichore lengths is provided in the supplementary material (Supplementary Data 1).

In our model, we chose the population *N* = 1000, the initial sequence length of 1024 bp (*L*_0_), the number of generations *m* = 1000, and used a 4-level (*ω* = 4) wavelet transformation. We repeated the experiment 100 times to ensure statistical robustness.

Our model also includes an upper threshold for the number of replication origins allowed in a genome (*Ori*_*max*_) to prevent an uncontrolled explosion in genome length. Genomes with a replication origin count greater than *Ori*_*max*_ are eliminated during the selection process. For bacteria, *Ori*_*max*_ is set to 1, for archaea, it is set to 5, while for eukaryotes, *Ori*_*max*_ is set to a value much greater than 1 (50 and 100). Although we use parameters such as *Ori*_*max*_, wavelet level *ω*, and mutation size 5% - 10% for computational convenience, the observed divergence between the bacterial and eukaryotic genome length is completely insensitive to the parameters listed above (Supplementary Data 2).

## Results

The variation in genome length (in bp) of single-origin and multi-origin sequences over 500 generations is shown in Fig. 5(a) and (b), respectively. The evolutionary minimization of the selection pressure *γ*, for single and multi-origin sequences, is shown in Fig. 5(c) and (d).

**Figure 5.**
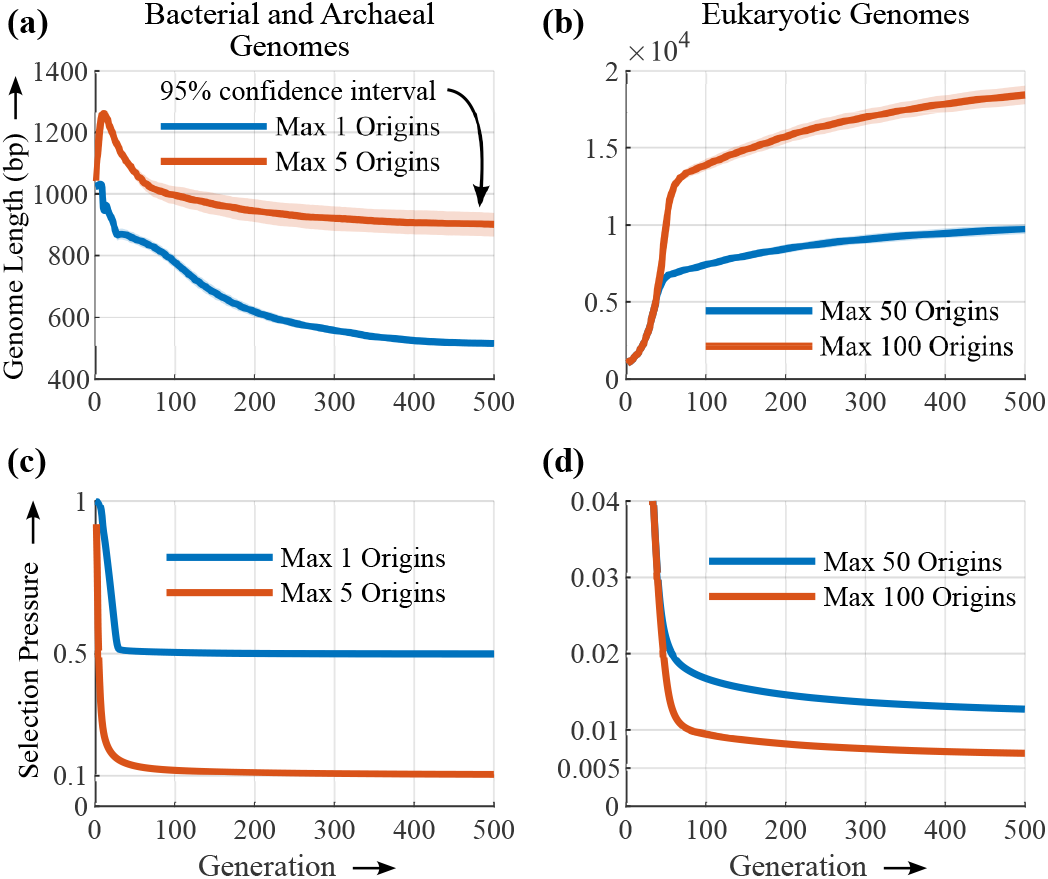
The evolution of genome length over generations. An initial population of 1000 purine-filled single-strand sequences is subjected to mutation and evolves under the selection pressure (*γ*) to minimize replication time while maximizing information storage capacity. The experiment is repeated 100 times, and the evolution of average genome length over 500 generations is shown, with a 95% confidence interval. (a) When each genome in the pool is constrained to have a single replication origin, mimicking bacterial genomes, the average genome length decreases over generations. Also shown is the genome length evolution when a few more origins are allowed, as in the case of archaea. (b) In contrast, when the sequences are allowed to accommodate many more replication origins, mimicking eukaryotic genomes, the average genome length increases. This is because the expansion in genome length of a multi-origin genome does not come at the expense of replication time. Unlike single-origin sequences, multi-origin sequences can parallelize replication across multiple replichores by replicating them independently and simultaneously, thereby reducing the replication time substantially, while maintaining a large genome length. Single-origin sequences cannot have more than two replichores, and hence cannot parallelize replication beyond these two, thus restricting their genome length. (c) and (d) Minimization of mutation pressure *γ*. For both single and multi-origin sequences, *γ* converges to 1*/*number of replichores. This convergence implies symmetrization of replichore lengths across a genome. The initial pool has no origins, and hence one replichore, and the initial value of *γ* is thus 1.

We observe that, in the absence of any restrictions on the number of replication origins, the genome tends to increase in length indefinitely (Fig. 5b). However, taking into account the scarcity of monomer resources and energy required to replicate longer genomes, we restrict genomes to have a maximum of *Ori*_*max*_ origins. When *Ori*_*max*_ is set high (*≫* 1), the average genome length is observed to increase over generations. On the other hand, when *Ori*_*max*_ is restricted to 1, mimicking a bacterial genome, the average length of the genome decreases over generations. Despite the applied selection pressure being identical in both scenarios, due to varying *Ori*_*max*_ constraints, single-origin genomes tend to lose sequence length, while multi-origin genomes tend to elongate their genomes. This behavior mirrors the evolutionary divergence between the lengths of the bacterial and eukaryotic genomes. In the context of the model, the explanation for this pattern of genome evolution is as follows.

Consider a genome with an asymmetric purine-pyrimidine composition, where all nucleotides at the 5′ end of the replication origin are pyrimidines, and those at the 3′ end are purines. The cumulative skew profile of this sequence forms a “V” shape, as illustrated in Fig. 6(a). Single-origin genomes with the two replichores emerging from their single origin exhibit such skew profiles. The single replication origin is located at the valley point of the “V”-shaped skew profile [69, 70, 71, 82]. The skew profile of a multi-origin genome, on the other hand, has multiple concatenated “V”-shapes, as shown in Fig. 6(b), where each arm of each “V” corresponds to a replichore, and the multiple valley points, to multiple origins. Mutations in the sequence alter this skew profile. The selection pressure *γ* depends on whether the genome undergoes deletion or insertion, and which replichore arm (shorter or longer) is affected due to the mutation. Fig. 6 shows a few skew profiles of the mutated sequences. Selection acts on the set of genomes with such altered skew profiles and prefers sequences that decrease the replication time and/or increase the information storage.

**Figure 6.**
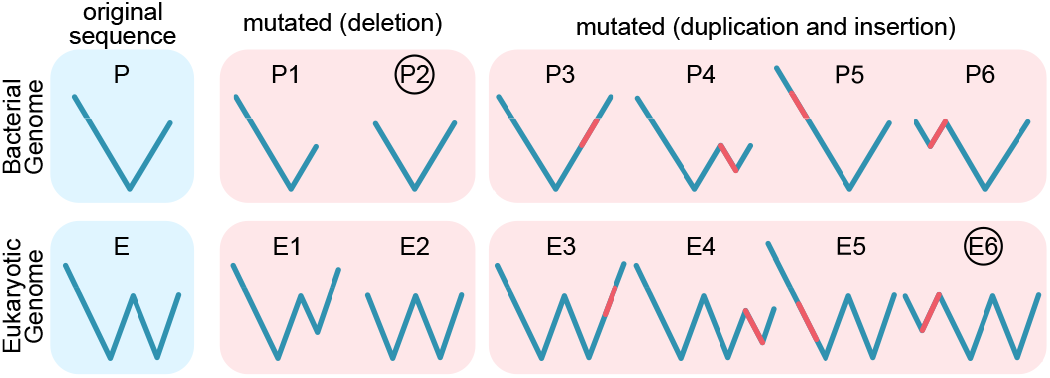
Effect of selection pressure on single- and multi-origin genomes. The left panel, in a blue background, shows the skew profiles of a single-origin and a multi-origin genome of an intermediate generation, before mutation. The right panel, in a red background, shows a few skew profiles of these genomes, after mutation. For multi-origin genomes, mutations adding an extra replichore originating from a new replication origin (e.g., ‘E6’) are preferred, as they increase information storage by elongating the genome without increasing the replication time (determined by the length of the longest replichore). In single-origin genomes, mutations that increase the length of the longest replichore (e.g., ‘P3’ and ‘P5’) or add a new replication origin (e.g., ‘P4’ and ‘P6’) are eliminated by selection pressure. Instead, mutations that shorten the longest replichore length, leading to a symmetrization of replichore lengths (e.g., ‘P2’), are favored, as they optimize the selection pressure (*γ*_opt_ = 1*/*2) by lowering the replication time.

In the single-origin genome (P), mutations involving elongations are not preferred by the selection, since it either increases the longest replichore length (e.g., P3 and P5), resulting in an increase in overall replication time, or adds an extra valley point to the sequence (e.g., P4 and P6), which is eliminated by the selection pressure, as bacteria are restricted to have a single replication origin. However, a mutation that shortens the longest replichore (e.g., P2) is selected because it decreases the replication time. Unlike single-origin genomes, in multi-origin genomes, our selection algorithm allows for the addition of more origins, and hence more replichores (e.g., E4 and E6). If these new replichores are not the longest among all replichores, mutated sequences containing them will be selected due to their increased information storage capacity and neutral influence on replication time (e.g., E6). Therefore, the size of multi-origin genomes continues to increase over generations through the incorporation of new origins and hence replichores, until selection restricts further increase in the number of origins due to the upper limit *Ori*_*max*_.

In both single- and multi-origin sequences, evolutionary selection pressures favor the symmetrization of replichore lengths. This phenomenon arises because replichores that are shorter than the longest replichore can undergo elongation through mutational processes, as this enhances the genome’s information storage capacity without increasing its replication time, thereby reducing *γ*. This aligns with the general observation of symmetric replichore lengths in bacteria [83, 84, 85]. Collectively, these selective forces drive two key outcomes: (1) the equilibration of replichore lengths and (2) the addition of one or more new origins, and hence replichores.

The observed genome length reduction in the single-origin genome (Fig. 5) is due to a bias of the selection pressure that favors deletions over insertions, *although both of these mutational events are equally probable* in our model. As explained in the previous paragraph, selection pressure favors the symmetrization of replichores. When two replichores of the single-origin sequence are of unequal length, symmetrization requires deletion of the longer replichore or insertion into the shorter replichore. Since our choice for the location of these two mutational events is entirely random and, therefore, evenly distributed over the lengths of the genome, the probability of choosing the shorter replichore for insertion or deletion will always be smaller than the probability of mutations occurring on the longer replichore. Although selection favors insertion into shorter replichore and deletion at longer replichore equally, since the latter is *stochastically more favored*, deletion occurs more frequently. This computational observation reproduces the strong evolutionary preference observed experimentally for deletions over insertions across domains [86, 11, 87, 88, 89, 90].

An initial increase in the average genome length is seen in Fig. 5. This is an artifact of our initial choice of genomes, each of which has been chosen as a homogeneous stretch of purines or pyrimidines. Therefore, these initial sequences have no origin, and the length of the entire genome is equal to the length of a replichore, with *γ* = 1. The deletions in these sequences do not alter the value of *γ*, since they affect both the numerator and the denominator of *γ* equally. However, duplications can reduce *γ* when the duplicating fragment is from the complementary strand, introducing a new replichore and an origin. The reduction in *γ* is due to the division of the genome into two replichores, due to the introduction of an origin, thereby reducing the replication time [62]. As a result, within our model, during the early stages of evolution, deletions are not favored by selection, and the average genome length increases.

### Symmetrization of replichore lengths leads to Chargaff’s Second Parity Rule

The evolution of genomic sequences in our model begins with a pool of sequences composed entirely of purines (R) on the Watson-strand and complementary pyrimidines (Y) on the Crick-strand. In successive generations, mutation involving the exchange of strand fragments between complementary strands introduces strand heterogeneity, that is, sequences with interspersed R and Y bases. As demonstrated in the earlier section, evolution under the defined selection pressure *γ* drives sequences to have replichores of equal length. Half of the replichores, positioned to the left of each origin, become pyrimidine-rich, while the remaining half, to the right of origins, are purine-rich. This symmetry in the cumulative nucleotide skew around replication origins [68] results in global parity in the purine and pyrimidine content throughout the genome, leading to Chargaff’s second parity rule (PR-2).

Chargaff’s first parity rule, identified before the discovery of DNA’s double-stranded structure [91, 92], revealed equal counts of adenine (A) and thymine (T), as well as guanine (G) and cytosine (C) in double-stranded DNA (dsDNA), a pattern now understood as a consequence of Watson-Crick base pairing. In contrast, Chargaff’s second parity rule (PR-2), which extends this symmetry to individual strands of dsDNA, lacks a universally accepted mechanistic explanation [93, 94, 95, 96]. Early hypotheses attributed PR-2 to adaptive intra-strand stem-loop formation, favoring local sequence inversions to achieve the functional benefits of self-complementarity [96, 97]. However, this rationale fails to account for PR-2’s prevalence in non-coding regions, where selective pressures for secondary structures are weak [98]. Alternative theories propose PR-2 as a manifestation of the law of large numbers or an emergent property of entropy maximization in large genomes, where stochastic shuffling of sequences via inversions and transpositions homogenizes base composition over time [99, 100, 101, 102, 103, 104]. However, these mechanisms rely on *no strand-bias* assumptions and do not account for an asymmetry in base substitution frequencies between purine and pyrimidine, i.e., *R → Y* ≠ *Y → R* [105, 106]. More importantly, while these theories account for global compositional symmetries, they neither explain nor reproduce local nucleotide compositional skews around replication origins that are prevalent in genomes of nearly all species [68, 69, 70, 71, 107], and are essential for incorporation of replication origins within Asymmetric Cooperativity model [63]. The near-universal presence of local violations of PR-2 points to their importance, specifically for replication origin functionality. Stochastic nucleotide shuffling would erase these local PR-2 violations as well, and hence will be counterproductive.

In our framework, although we incorporate inter-strand shuffling, PR-2 emerges not as a passive outcome of random sequence shuffling but as an adaptive response to selection pressure (Fig. 7). Here, selection pressure eliminates any bias in the base substitution frequencies between purines and pyrimidines. Any alteration in the symmetric, V-shaped profile of the cumulative nucleotide skew (see Fig. 6) due to the bias in base substitution frequencies is not tolerated by the selection, since it would adversely affect the replication time, by making the replichore lengths asymmetric. Therefore, selection acts against such biased substitutions, restoring the symmetry of the cumulative skew diagram and that of the replichore lengths, and hence the global equivalence in the purine-pyrimidine content of a DNA single-strand. The evolutionary trajectory of purine content in our simulations (Fig. 7) illustrates this dynamic, demonstrating a rapid convergence to R/Y parity concurrent with replichore symmetrization. Note that, although the mean R/Y composition equilibrates to 50% in both selective and neutral evolutionary processes (red and blue curves in Fig. 7), the variance in R/Y composition is larger for neutral evolution, and the equilibration takes longer [104], when compared with the evolution under selective pressure. This suggests that strong selective pressure leads to strict compliance with PR-2, whereas non-adaptive evolution allows for deviations from PR-2. If the selection pressure for short replication time is weak or non-existent, as in the case of mitochondria, plasmids and viroids, the PR-2 compliance requirement vanishes, according to our model. There is a lack of need to maintain symmetric replichore lengths to minimize replication time in these genomes, since the rate-limiting step for their replication is the replication time of the host genomes, which are much larger. This prediction has been validated experimentally [94, 108]. Chloroplasts’ replication is independent of their host cell cycle [109], and it is, therefore, PR-2 compliant.

**Figure 7.**
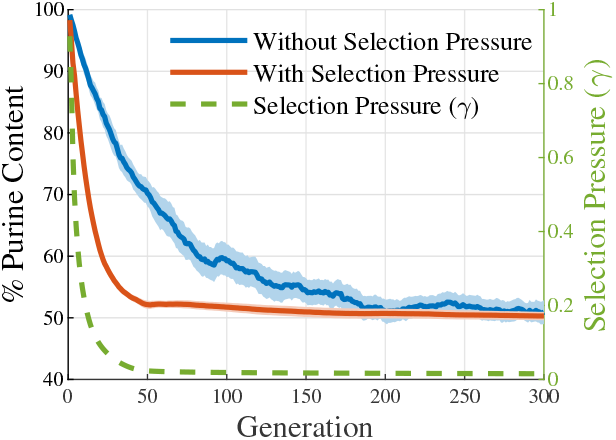
Evolutionary trajectories of base composition over generations. An initial pool of double-stranded DNA, with the Watson strand composed entirely of purines (R) and the Crick strand of pyrimidines (Y), evolves through mutations (inversions and inverted transpositions). The variation in purine content of the Watson strand over generations is shown, with shaded regions indicating 95% confidence interval. **Blue curve:** Strand evolution without any selection pressure, exhibiting gradual R/Y parity in DNA single-strand, due to stochastic inter-strand shuffling. This demonstrates PR-2 emerging from stochasticity alone. These sequences destroy any structured nucleotide skew profiles required for replication origins, and observed in most genomes. **Red curve:** Under selection, R/Y parity emerges more rapidly via replichore symmetrization, which is a direct consequence of the selection pressure to minimize replication time by balancing replichore lengths. Selection maintains the nucleotide skew profiles around replication origins, aligning with experimental observations. Selection also preserves PR-2 by eliminating mutational biases for specific nucleotides, removing the necessity for the no-strand bias assumption, used in explanations based on neutral processes.

### Influence of endosymbiont power supply on genome length

According to our model above, bacteria and archaea tend to minimize their genome length due to the limitation of a single (or a few) replication origin, whereas eukaryotes tend to acquire genome content because of their ability to accommodate multiple replication origins. The rationale for our choice to restrict the number of origins of bacteria to one, while allowing the eukaryotes to have many more, is to align the model with observations. A deeper reason for this choice lies in the bioenergetic requirement for maintaining multiple replication origins, which is to provide activated monomers and an energy supply for the replication machinery *simultaneously* to multiple replicating segments of a large genome. This requirement is met through the endosymbiotic relationship of eukaryotes with mitochondria or chloroplasts [9, 17].

To test this hypothesis, we correlated nuclear genome content with these organelle counts using data from Jordan G. Okie et al.[110] and other literature sources. As the energetic cost of genome maintenance depends on total genomic content, we used the product of 1C genome size and ploidy level, rather than 1C alone. Both ordinary least squares (OLS) and phylogenetic generalized least squares (PGLS) regressions were performed (Fig. 8), with PGLS applied only to species with phylogenetic data from TimeTree [111]. A weak phylogenetic signal (low *K* values; see Table 1) was observed in both traits, nuclear genome content and the organelle count, of the mitochondria data set, suggesting that PGLS is not essential for this [112]. The low phylogenetic signal in mitochondrial count likely reflects its variability across the cell cycle, while in genome content, it may be due to the C-value paradox. The ordinary linear regression correlation between organelle abundance and genome content suggests that genome length in eukaryotes is constrained by the energy required to replicate multiple regions simultaneously, supplied by mitochondria and/or chloroplasts.

**Figure 8.**
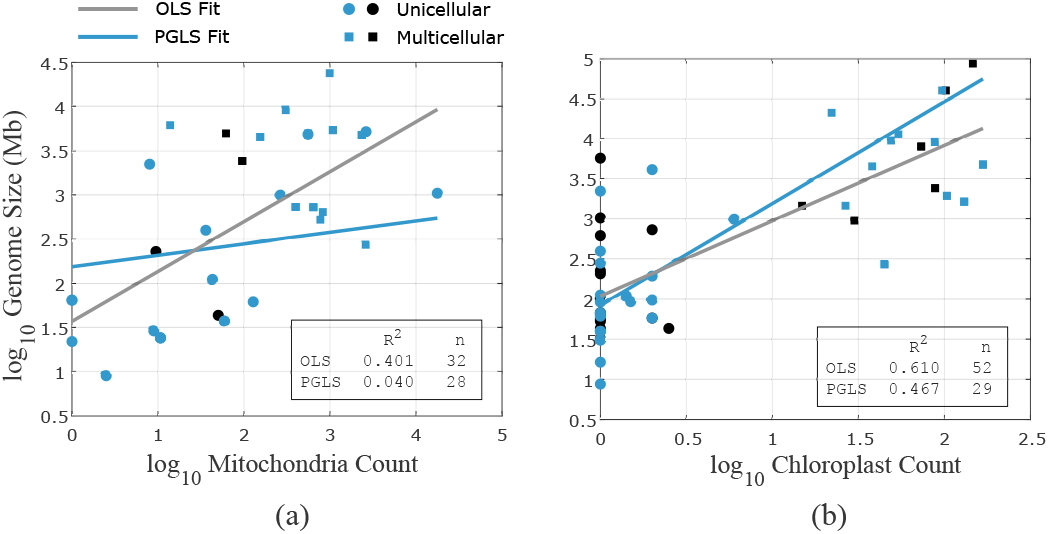
Correlation between organelle count and nuclear genome content. (a) The correlation between the number of mitochondria and nuclear genome content in 32 eukaryotic cells, out of which 28 species have phylogenetic data available, and their phylogenetic generalized least squares regression is done. Note that the phylogenetic signal in both traits (mitochondria count and genome size) is low (*K <*.5), and hence the PGLS correlation coefficient *R*^2^ is not significant. (b) Correlation between the number of chloroplasts and genome length in 52 eukaryotic cells, out of which 29 are considered in PGLS analysis, whose phylogenetic data were available. The organelle count data is provided by Jordan G. Okie et al. [110]. The nuclear genome content is taken from the literature, where the genome size (1C) is multiplied by the ploidy level of the species. The genome length is positively correlated with mitochondrial or chloroplast count, supporting the argument that the replication of longer genomes of eukaryotes is carried out with the energy provided by the organelles.

This limitation imposed on genome length by cellular energetics is taken into account in our model by limiting the number of origins to a set value, *Ori*_*max*_. In bacteria and archaea, which lack mitochondria/chloroplasts, the number of origins is generally limited to 1 (or a few), and therefore we take *Ori*_*max*_ = 1 or 5. In eukaryotes, the number of origins can be of the order of tens of thousands, and is limited only by power supply availability, as argued above (8). We therefore set *Ori*_*max*_ to 50 or 100, for computational convenience, to explore the genome length divergence between bacteria and eukaryotes.

### Upper and lower bound of genome length

As explained before, the selection pressure symmetrizes the replichore length and adds new origins, and thus replichores, to maximize information storage capacity. After evolving the sequences for a considerable number of generations, the selection produces genomes with nearly equal replichore lengths and the maximum number of origins (*Ori*_*max*_ = 1 for single-origin and *Ori*_*max*_ *>>* 1 for multi-origin), thus minimizing the selection pressure. The theoretical optimum genome length under these conditions is

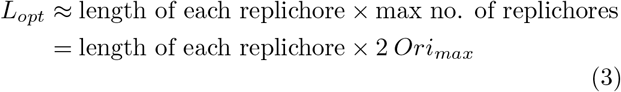

Therefore, the optimum selection pressure is

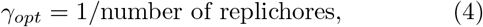

The selection pressure can be seen to converge to this optimum in Fig. 5(c) & (d). For single-origin genome, *γ*_*opt*_ = ^1^, since a single origin can only support two replichores. Whereas, for multi-origin genomes, *γ*_*opt*_ is 0.01 and 0.005, since, at optimum genome length, they will have *≈* 100 and *≈* 200 replichores, corresponding to 50 and 100 origins, respectively.

#### Lower bound

The model introduces a wavelet-based resolution limit for distinguishing replication origins. Following a *w*-level wavelet transform of the genome, pairs of peaks or valleys in the nucleotide skew profile separated by fewer than 2^*w*^ nucleotides become indistinguishable, as the transform disregards small-scale variations. This imposes a minimum threshold of 2^*w*^ base pairs (bp) on replichore lengths. This threshold may serve as a proxy for the minimal genomic information necessary for cellular viability, reflecting evolutionary constraints on the smallest genome length. Consequently, in our simulations, the smallest size achievable for a single-origin genome is of the order of 2 *×* 2^*w*^ nucleotides. Within the model, this lower bound reflects the computational resolution limit for identification of origins, whereas, in the evolutionary dynamics captured by the model, this lower bound reflects the need for sufficient informational complexity for cellular maintenance and replication machinery [20, 113, 114, 115, 116]. Our choice of a specific wavelet level ensures that the genome length does not reduce below a certain viability limit.

#### Upper bound

One can make an interesting observation from the eq. 4: The optimum length of a eukaryotic genome does not depend on the length of the replichores; only on their number. A genome can therefore increase its own length by increasing the length of each of its replichores evenly, without influencing *γ*. Such an alteration increases both the replication time and the information storage capacity equally, thereby nullifying its effect on the selection pressure *γ*. However, our stochastic model cannot make such concerted changes at multiple locations in the genome, and hence cannot alter the length of the genome this way. When the selection pressure is low, evolution, on the other hand, can alter the genome length by progressively lengthening each of the replichores, by temporarily tolerating an uneven distribution of replichore lengths. This would result in very different genome lengths and corresponding variance in replication times, due to variation in replichore length, even within very similar eukaryotic species, as has been observed abundantly [23, 24, 117, 118]. Therefore, our model does not impose a strict upper limit on the genome length, although it converges to a constant genome length for a fixed number of origins, *Ori*_*max*_. The evolutionarily optimized eukaryotic genome length for a constant *Ori*_*max*_ value depends on the average length of the initial genome pool, which can be altered to produce a longer or shorter optimized genome.

### Large variance of eukaryotic genome length due to low selection pressure for replication time

In order to verify our statement above that the genome length can vary drastically even within similar species when the selection pressure for replication time is low, we reduce the cost of replication time in our evaluation of selection pressure, by raising it to a power *α*, where 0 *≤ α ≤* 1. The modified expression for selection pressure reads

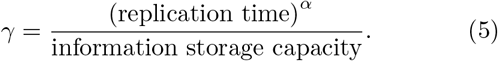

This modification allows for uneven distribution of replichore lengths, and the algorithm tolerates an increase in the replication time by prioritizing information storage capacity, thus enlarging the genome, as explained above. When the importance of replication time reduces, and its cost (*α*) goes down, the variance in the lengths of the genomes of our initial population increases with generation, as seen from the widening of the confidence interval with generation in Fig. 9.

**Figure 9.**
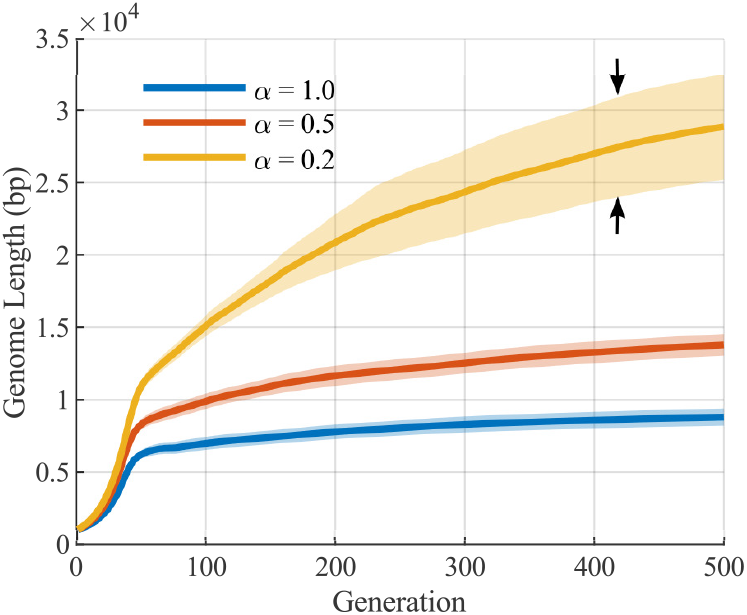
Genome length evolution under reduced selection pressure for replication time. In our model, the replication time is dictated by the length of the longest replichore of the genome. Under low selection pressure for replication time (i.e., low *α*), replichores expand to enhance information storage capacity while minimally impacting the cost of replication time. This drives progressive genome elongation over generations. This reduced pressure also permits uneven replichore lengths across the genome, as opposed to the optimized symmetrical replichore lengths observed under strong selection. Consequently, genome length variability within a set of related species increases over time, as reflected in the widening confidence intervals for low *α*. This explains the genome length variability experimentally observed across species and within conspecific organisms. The confidence interval shown here is calculated with 30 samples, for the purpose of visual demonstration. The increase in variance with the reduction in *α* persists for higher sample numbers.

**Figure 10.**
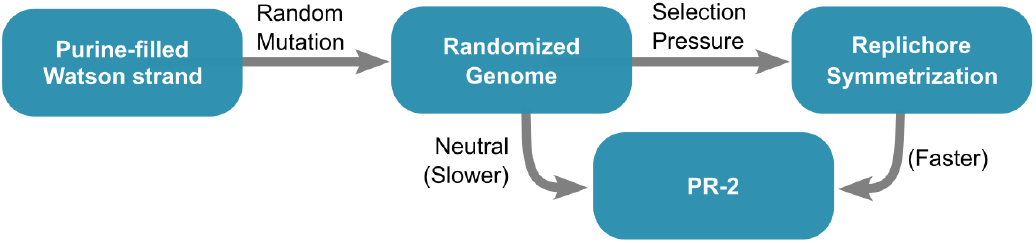
Neutral and adaptive evolutionary trajectories leading to Chargaff’s second parity rule (PR-2). In the neutral process, starting from a purine-filled Watson strand of low fitness (high *γ*), genomes may reach a state of equal purine (R) and pyrimidine (Y) content on a single strand, i.e. PR-2, solely through a large number of mutations (inversions and inverted transpositions), occurring neutrally without providing any adaptive benefit. The rate of convergence to PR-2 through this route is slower, and the PR-2 compliance is less strict. Moreover, the nucleotide skews that dictate the V-shaped replichore structure are averaged out, nullifying the sequence signature corresponding to replication origins. Alternatively, as demonstrated in this work, selection pressure for rapid replication can *swiftly enforce* PR-2 through replichore length symmetrization, resulting in a *strict* compliance with this parity rule. This adaptive process also retains the sequence signature that determines replication origins.

Such computationally observed significant variation in genome length has been documented across species [119, 120, 121] and even within conspecific populations [57, 117, 118, 122], an observation whose evolutionary mechanism is deeply contested. This constitutes the central mystery of the C-value paradox, a set of observations of uncertain evolutionary origins [22, 24]. Prevailing hypotheses posit that persistent upward mutation pressure drives C-value (a measure of genome content) expansion, with species exhibiting slower cellular division rates being more tolerant of random DNA accumulation [23, 55, 123]. This framework is supported by evidence demonstrating strong negative correlations between genome length and both mitotic and meiotic division rates [58, 59]. Our computational model provides an explanation for the above observation, where low selection pressure for replication time shifts the evolutionary trajectory towards maximizing information storage capacity at the expense of replication time. Apart from increasing the genome content over generations (C-value), this also increases the variance of the genome length, leading to drastically different genome lengths even within conspecific organisms (Fig. 9). Whether the accumulated genome carries information or not cannot be answered within our model, another mystery of the C-value paradox.

## Discussion

The divergence between the lengths of genomes of bacteria, archaea, and eukaryotes has been an enduring enigma, noted since the first systematic C-value measurements carried out in the 1950s. The nearly 100-fold difference in cell sizes between bacteria and eukaryotes, and the huge increase in the structural and functional complexity of eukaryotes, are generally attributed to this difference. The purpose of this paper is to investigate the evolutionary origin of this divergence. Here, we identify replication time and information storage capacity as the two primary determinants of genome size, with the former influenced by nucleotide skews, and the latter strongly constrained by cellular energetics. The interplay between these two variables dictates the evolutionary trajectory of genome size, for any given species.

We quantify this interplay between replication time and information storage capacity by defining Selection Pressure simply as the ratio of the above two variables. Evolution acts on the genome, attempting to minimize the replication time while maximizing the information storage capacity. By rewriting the selection pressure in terms of the length of the replichores and the total size of the genome, we make the selection pressure computable. We model the evolution of a pool of genomes by introducing stochastic mutations, randomly adding or deleting a fraction of the total genome, and evaluating the effect of the selection pressure on the mutated genomes.

We observe that bacterial genomes, modeled here as genomes with a *single* replication origin, lose genome size with evolving generations. This is due to the inability of bacterial genomes to parallelize their replication by dividing the genome into multiple segments (replichores) that can replicate independently and simultaneously, because of the restriction on the number of origins. This restriction reduces the genome size for bacteria, since only two simultaneously replicating segments are allowed, and any increase in the size of these segments increases the replication time, and is thus evolutionarily disfavored. On the other hand, we observe that eukaryotic genomes tend to increase their genome size indefinitely, without incurring a cost from increased replication time, since the model’s allowance of a large number of origins for eukaryotic genomes allows for massive parallelization of genome replication, by dividing the genome into multiple independently and simultaneously replicating segments. This indefinite expansion of the eukaryotic genome is curtailed only by cellular energetic constraints, the need for activated monomers, and energy supply for simultaneously replicating thousands of genomic segments. This constraint is experimentally demonstrated by the linear relationship between the number of organelles and genome size in eukaryotes (Fig. 8). We model this constraint by limiting the number of origins allowed for eukaryotic genomes.

The inclusion of the influence of purine–pyrimidine compositional skew on replication rate enables the model to reproduce several experimentally observed phenomena, beyond explaining genome size divergence across domains, such as (a) reproduction of Chargaff’s second parity rule (PR-2) compliance. (b) A general preference for deletion mutations over insertions (Fig. 6a) [11, 86, 87, 88, 89, 90]. (c) A tendency to equalize the lengths of all replichores of the genome, as indicated by the optimized *γ* value (Fig. 5c,d) [46, 83, 84, 85, 124] (d) Correlation between genome size and the number of mitochondria/chloroplasts (Fig. 8) [110, 125] (e) Increase in size [58, 59] and the variance of genome size resulting from a reduced selection pressure for replication time minimization (C-value paradox) (Fig. 9) [57, 117, 118, 122].

Our model demonstrates that Chargaff’s second parity rule (PR-2), i.e., the symmetry of purine (R) and pyrimidine (Y) frequencies within individual DNA strands, emerges as a direct consequence of selection for replication efficiency. When initialized with purine-filled single-strand sequences, evolution under the defined selection pressures drives them toward parity, yielding strands with equal purine and pyrimidine content. Critically, this symmetry is not a natural outcome of stochastic inter-strand sequence shuffling or thermodynamic entropy maximization, but an adaptive response to selection for balanced replichore lengths. Any compositional bias in a strand would generate asymmetrical replichores, delaying replication and reducing fitness. Thus, PR-2 in our framework reflects an evolutionary optimization: the equalization of R/Y content is a byproduct of selection to harmonize replichore architecture, ensuring efficient bidirectional replication. These results suggest that PR-2 is a signature of replication-driven adaptation, rather than an outcome of neutral processes.

Our model can be tested using *in-vitro* evolution experiments on self-replicating DNA sequences that involve deletions and insertion mutations. Under strong selection pressure for faster replication, the sequences should evolve towards a replichore structure with a central origin of replication and equal and opposite nucleotide skews on the two replichore arms. However, when primed to replicate from the ends, these sequences should evolve toward purine/pyrimidine-filled single strands. When the supply of monomers is adequate, the more fit sequences would exhibit multiple origins. Any imbalance in the lengths of the two replichores of the evolutionarily superior sequences should adversely affect the sequence’s fitness.

Our explanation for the divergence between bacterial, archaeal, and eukaryotic genome size rests primarily on the number of origins used during DNA replication: using more origins reduces the replication time, all else being equal. Multicellular eukaryotes appear to have invented a new degree of freedom to modulate their cell replication time depending on tissue-level spatial, temporal, energetic, and environmental constraints by employing an appropriate number of origins. As has been amply demonstrated, multicellular organisms do not utilize all available origins of replication to replicate their DNA. Only about 30% of the origins of the human genome are constitutively fired, with the utilization of the rest depending on the tissue/organism-level requirements [126, 127, 128]. This top-down control of replication origin firing partly enables these organisms to maintain specialized organs with disparate cell-cycle rates, such as human skin and colon, where rapid cell-cycle rates are crucial, and neurons in the brain, which rarely replicate, presumably to preserve information [129, 130, 131, 132]. An important sequence characteristic that segregates multicellular eukaryotic origins into constitutive, latent, and dormant sets is the magnitude of nucleotide skew at the origin locations [74]. Our model above also uses these nucleotide skews to identify the locations of origins, although the magnitude of the skew is not utilized for determining the efficiency of origin firing, a simplification that will be removed in a later article. We speculate that the local loss of such top-down control on replication origin selection in various tissues leads to rapid replication and, consequently, carcinogenesis [133]. Regulation of the number and firing efficiency of replication origins can modulate genome architecture at evolutionary timescales, while organismal top-down control appears to tame the origins into serving the individual organism at the timescale of the lifetime of that individual.

## Limitations of the model

Our model relies on simplified, albeit biologically motivated, assumptions and omits several factors that could significantly influence genome size evolution. For instance, while DNA replication across replichores and chromosomes is known to be semi-parallel and temporally ordered (Fig. 3), our model assumes that all replichores are simultaneously replicated across the entire genome. Temporal ordering cannot be captured within our model without identifying the constraints, such as nutrient availability, on the organism that leads to such partial serialization of replication. The model doesn’t include the impact of nucleotide skew on replication fork speed, which we have reported in our observation (Fig. 4(d). Relating the magnitude of skew to the replication fork speed in our model would lead to the erroneous conclusion that the fork speed would progressively reduce as an organism evolves, since the nucleotide skew magnitudes get progressively lower as the simulation proceeds. Inclusion of a countervailing evolutionary force against this fork speed reduction is reserved for a future article. Moreover, although we talked about origin firing efficiency and its relationship to the magnitude of nucleotide skews, we have not incorporated them into the current model. Therefore, our model is currently unable to distinguish between constitutive, latent and dormant origins.

The model does not allow for ploidy, as it considers total nuclear DNA as a single sequence and does not allow for whole-genome duplication during the mutation, which would lead to ploidy variations. The model is incapable of distinguishing between coding and non-coding DNA, implicitly treating any genome expansion as adaptive, a point of ongoing debate. Furthermore, ecological factors such as population size, nutrient availability, and other environmental pressures are also not included. Finally, the model does not fully capture archaeal genome dynamics: despite possessing more replication origins than bacteria, archaeal genomes remain similar in size, deviating from the model’s predictions. Incorporating these factors in future work could improve predictive accuracy and better reflect the diversity of genomic patterns observed in nature.

## Acknowledgments

Support for this work was provided by the Science & Engineering Research Board (SERB), Department of Science and Technology (DST), India, through a Core Research Grant with file no. CRG/2020/003555 and a MATRICS grant with file no. MTR/2022/000086.

## Supplementary Information

Details of the algorithms and parameter settings used to generate the plots are provided in Supplementary Information 1.

Additional simulation outputs generated with varying parameter sets, as well as the phylogenetic tree used in this study, are provided in Supplementary Information 2. The MATLAB code for simulating genome evolution, along with the datasets used in the analyses, are publicly available at the following GitHub repository: https://github.com/ParthaTbio/Genome_Evolution.

## References

[1] T Ryan Gregory. “Understanding natural selection: essential concepts and common misconceptions”. In: Evolution: Education and outreach 2 (2009), pp. 156–175.

[2] Freeman Dyson. Origins of life. Cambridge University Press, 1999.

[3] Radhey S Gupta and G Brian Golding. “The origin of the eukaryotic cell”. In: Trends in biochemical sciences 21.5 (1996), pp. 166–171.

[4] T Martin Embley and William Martin. “Eukaryotic evolution, changes and challenges”. In: Nature 440.7084 (2006), pp. 623–630.

[5] William F Martin, Sriram Garg, and Verena Zimorski. “Endosymbiotic theories for eukaryote origin”. In: Philosophical Transactions of the Royal Society B: Biological Sciences 370.1678 (2015), p. 20140330.

[6] Itamar Sela, Yuri I Wolf, and Eugene V Koonin. “Theory of prokaryotic genome evolution”. In: Proceedings of the National Academy of Sciences 113.41 (2016), pp. 11399–11407.

[7] Michael Lynch and John S Conery. “The origins of genome complexity”. In: science 302.5649 (2003), pp. 1401–1404.

[8] Eduardo PC Rocha. “The organization of the bacterial genome”. In: Annual review of genetics 42.1 (2008), pp. 211–233.

[9] Nick Lane. “Energetics and genetics across the prokaryote-eukaryote divide”. In: Biology direct 6 (2011), pp. 1–31.

[10] Katsumi Chiyomaru and Kazuhiro Takemoto. “Revisiting the hypothesis of an energetic barrier to genome complexity between eukaryotes and prokaryotes”. In: Royal Society open science 7.2 (2020), p. 191859.

[11] Alex Mira, Howard Ochman, and Nancy A Moran. “Deletional bias and the evolution of bacterial genomes”. In: TRENDS in Genetics 17.10 (2001), pp. 589–596.

[12] Piotr Bentkowski, Cock Van Oosterhout, and Thomas Mock. “A model of genome size evolution for prokaryotes in stable and fluctuating environments”. In: Genome biology and evolution 7.8 (2015), pp. 2344– 2351.

[13] Carolina A Martinez-Gutierrez and Frank O Aylward. “Genome size distributions in bacteria and archaea are strongly linked to evolutionary history at broad phylogenetic scales”. In: PLoS Genetics 18.5 (2022), e1010220.

[14] Alejandro Rodríguez-Gijón et al. “Linking prokaryotic genome size variation to metabolic potential and environment”. In: ISME communications 3.1 (2023), p. 25.

[15] Stephan Fischer et al. “A model for genome size evolution”. In: Bulletin of mathematical biology 76 (2014), pp. 2249–2291.

[16] Marco Colnaghi, Nick Lane, and Andrew Pomiankowski. “Genome expansion in early eukaryotes drove the transition from lateral gene transfer to meiotic sex”. In: Elife 9 (2020), e58873.

[17] Nick Lane and William Martin. “The energetics of genome complexity”. In: Nature 467.7318 (2010), pp. 929–934.

[18] Yubo Hou and Senjie Lin. “Distinct gene number-genome size relationships for eukaryotes and non-eukaryotes: gene content estimation for dinoflagellate genomes”. In: PloS one 4.9 (2009), e6978.

[19] Dan Graur and Wen-Hsiung Li. Molecular evolution. Sinauer Associates, Sunderland, MA, 1997.

[20] Alexander V Markov, Valery A Anisimov, and Andrey V Korotayev. “Relationship between genome size and organismal complexity in the lineage leading from prokaryotes to mammals”. In: Paleontological Journal 44 (2010), pp. 363–373.

[21] Jr Thomas CA. “The genetic organization of chromosomes.” In: (1971).

[22] Susumu Ohno. “So much “junk” DNA in our genome. In “Evolution of Genetic Systems”“. In: Brookhaven symposium in biology. Vol. 23. 1972, pp. 366–370.

[23] Mark Pagel and Rufus A Johnstone. “Variation across species in the size of the nuclear genome supports the junk-DNA explanation for the C-value paradox”. In: Proceedings of the Royal Society of London. Series B: Biological Sciences 249.1325 (1992), pp. 119–124.

[24] T Ryan Gregory. “Coincidence, coevolution, or causation? DNA content, cell size, and the C-value enigma”. In: Biological reviews 76.1 (2001), pp. 65–101.

[25] T Cavalier-Smith. “Skeletal DNA and the evolution of genome size”. In: Annual review of biophysics and bioengineering 11.1 (1982), pp. 273–302.

[26] Eörs Szathmáry and John Maynard Smith. The major transitions in evolution. WH Freeman Spektrum Oxford, 1995.

[27] Charles A Knight, Nicole A Molinari, and Dmitri A Petrov. “The large genome constraint hypothesis: evolution, ecology and phenotype”. In: Annals of botany 95.1 (2005), pp. 177–190.

[28] Stephen Nayfach and Katherine S Pollard. “Average genome size estimation improves comparative metagenomics and sheds light on the functional ecology of the human microbiome”. In: Genome biology 16.1 (2015), p. 51.

[29] Nurhani Mat Razali, Boon Huat Cheah, and Kalaivani Nadarajah. “Transposable elements adaptive role in genome plasticity, pathogenicity and evolution in fungal phytopathogens”. In: International journal of molecular sciences 20.14 (2019), p. 3597.

[30] Craig F Barrett et al. “Ancient polyploidy and genome evolution in palms”. In: Genome Biology and Evolution 11.5 (2019), pp. 1501–1511.

[31] Lilian-Lee B Müller, Gerhard Zotz, and Dirk C Albach. “Bromeliaceae subfamilies show divergent trends of genome size evolution”. In: Scientific Reports 9.1 (2019), p. 5136.

[32] Claus-Peter Stelzer, Maria Pichler, and Anita Hatheuer. “Linking genome size variation to population phenotypic variation within the rotifer, Brachionus asplanchnoidis”. In: Communications Biology 4.1 (2021), p. 596.

[33] Zhao Chen et al. “Altitudinal patterns in adaptive evolution of genome size and inter-genome hybridization between three Elymus species from the Qinghai– Tibetan plateau”. In: Frontiers in Ecology and Evolution 10 (2022), p. 923967.

[34] Jesper Boman and Göran Arnqvist. “Larger genomes show improved buffering of adult fitness against environmental stress in seed beetles”. In: Biology Letters 19.1 (2023), p. 20220450.

[35] Lucas Serra Moncadas et al. “Freshwater genome-reduced bacteria exhibit pervasive episodes of adaptive stasis”. In: nature communications 15.1 (2024), p. 3421.

[36] Thomas Cavalier-Smith. “Economy, speed and size matter: evolutionary forces driving nuclear genome miniaturization and expansion”. In: Annals of botany 95.1 (2005), pp. 147–175.

[37] Hyunmin Lee, Zhaolei Zhang, and Henry M Krause. “Long noncoding RNAs and repetitive elements: junk or intimate evolutionary partners?” In: TRENDS in Genetics 35.12 (2019), pp. 892–902.

[38] Evgeniy S Balakirev and Francisco J Ayala. “Pseudogenes: are they “junk” or functional DNA?” In: Annual review of genetics 37.1 (2003), pp. 123–151.

[39] Nils G Walter. “Are non-protein coding RNAs junk or treasure? An attempt to explain and reconcile opposing viewpoints of whether the human genome is mostly transcribed into non-functional or functional RNAs”. In: BioEssays 46.4 (2024), p. 2300201.

[40] Peter Andolfatto. “Adaptive evolution of non-coding DNA in Drosophila”. In: Nature 437.7062 (2005), pp. 1149–1152.

[41] Aurélie Hua-Van et al. “The struggle for life of the genome’s selfish architects”. In: Biology direct 6.1 (2011), p. 19.

[42] Jeffrey Snowbarger, Praveen Koganti, and Charles Spruck. “Evolution of repetitive elements, their roles in homeostasis and human disease, and potential therapeutic applications”. In: Biomolecules 14.10 (2024), p. 1250.

[43] Julie Blommaert. “Genome size evolution: towards new model systems for old questions”. In: Proceedings of the Royal Society B 287.1933 (2020), p. 20201441.

[44] Patrick Alfred Pierce Moran. “Random processes in genetics”. In: Mathematical proceedings of the cambridge philosophical society. Vol. 54. 1. Cambridge University Press. 1958, pp. 60–71.

[45] Yuri I Wolf et al. “The universal distribution of evolutionary rates of genes and distinct characteristics of eukaryotic genes of different apparent ages”. In: Proceedings of the National Academy of Sciences 106.18 (2009), pp. 7273–7280.

[46] Guenter Albrecht-Buehler. “Inversions and inverted transpositions as the basis for an almost universal “format” of genome sequences”. In: Genomics 90.3 (2007), pp. 297–305.

[47] Saugat Bolakhe. “Noncoding RNAs have a key role in butterfly speciation. What about other flora and fauna?” In: Proceedings of the National Academy of Sciences 122.28 (2025), e2515930122.

[48] Rita Rebollo, Mark T Romanish, and Dixie L Mager. “Transposable elements: an abundant and natural source of regulatory sequences for host genes”. In: Annual review of genetics 46.1 (2012), pp. 21–42.

[49] Flávio SJ de Souza, Lucía F Franchini, and Marcelo Rubinstein. “Exaptation of transposable elements into novel cis-regulatory elements: is the evidence always strong?” In: Molecular biology and evolution 30.6 (2013), pp. 1239–1251.

[50] Corinne N Simonti, Mihaela Pavličev, and John A Capra. “Transposable element exaptation into regulatory regions is rare, influenced by evolutionary age, and subject to pleiotropic constraints”. In: Molecular Biology and Evolution 34.11 (2017), pp. 2856–2869.

[51] Erica Gasparotto et al. “Transposable elements cooption in genome evolution and gene regulation”. In: International Journal of Molecular Sciences 24.3 (2023), p. 2610.

[52] Jorge Ruiz-Orera et al. “Origins of de novo genes in human and chimpanzee”. In: PLoS genetics 11.12 (2015), e1005721.

[53] Dan I Andersson, Jon Jerlström-Hultqvist, and Joakim Näsvall. “Evolution of new functions de novo and from preexisting genes”. In: Cold Spring Harbor perspectives in biology 7.6 (2015), a017996.

[54] Henrik Kaessmann. “Origins, evolution, and phenotypic impact of new genes”. In: Genome research 20.10 (2010), pp. 1313–1326.

[55] Michael David Bennett. “Nuclear DNA content and minimum generation time in herbaceous plants”. In: Proceedings of the Royal Society of London. Series B. Biological Sciences 181.1063 (1972), pp. 109–135.

[56] VB Ivanov. “DNA content in the nucleus and rate of development in plants”. In: Soviet Journal of Developmental Biology 9 (1978), pp. 39–53.

[57] MA Mowforth and JP Grime. “Intra-population variation in nuclear DNA amount, cell size and growth rate in Poa annua L.” In: Functional Ecology (1989), pp. 289– 295.

[58] Irena Šímová and Tomáš Herben. “Geometrical constraints in the scaling relationships between genome size, cell size and cell cycle length in herbaceous plants”. In: Proceedings of the Royal Society B: Biological Sciences 279.1730 (2012), pp. 867–875.

[59] Dennis Francis, M Stuart Davies, and Peter W Barlow. “A strong nucleotypic effect on the cell cycle regardless of ploidy level”. In: Annals of Botany 101.6 (2008), pp. 747–757.

[60] Maud I Tenaillon et al. “Testing the link between genome size and growth rate in maize”. In: PeerJ 4 (2016), e2408.

[61] Avraham Greenberg and Itamar Simon. “S phase duration is determined by local rate and global organization of replication”. In: Biology 11.5 (2022), p. 718.

[62] Hemachander Subramanian and Robert A Gatenby. “Evolutionary advantage of anti-parallel strand orientation of duplex DNA”. In: Scientific Reports 10.1 (2020), p. 9883.

[63] Parthasarathi Sahu et al. “High Nucleotide Skew Palindromic DNA Sequences Function as Potential Replication Origins due to their Unzipping Propensity”. In: Journal of Molecular Evolution (2024), pp. 1–15.

[64] Dominik Boos and Pedro Ferreira. “Origin firing regulations to control genome replication timing”. In: Genes 10.3 (2019), p. 199.

[65] Prasanta K Patel et al. “DNA replication origins fire stochastically in fission yeast”. In: Molecular biology of the cell 17.1 (2006), pp. 308–316.

[66] Xindan Wang et al. “Replication and segregation of an Escherichia coli chromosome with two replication origins”. In: Proceedings of the National Academy of Sciences 108.26 (2011), E243–E250.

[67] Rivka Rudner, John D Karkas, and Erwin Chargaff. “Separation of B. subtilis DNA into complementary strands. 3. Direct analysis.” In: Proceedings of the National Academy of Sciences 60.3 (1968), pp. 921–922.

[68] Jean R Lobry. “Asymmetric substitution patterns in the two DNA strands of bacteria.” In: Molecular biology and evolution 13.5 (1996), pp. 660–665.

[69] Andrei Grigoriev. “Analyzing genomes with cumulative skew diagrams”. In: Nucleic acids research 26.10 (1998), pp. 2286–2290.

[70] Elisabeth RM Tillier and Richard A Collins. “The contributions of replication orientation, gene direction, and signal sequences to base-composition asymmetries in bacterial genomes”. In: Journal of Molecular Evolution 50 (2000), pp. 249–257.

[71] Michael J McLean, Kenneth H Wolfe, and Kevin M Devine. “Base composition skews, replication orientation, and gene orientation in 12 prokaryote genomes”. In: Journal of molecular evolution 47 (1998), pp. 691– 696.

[72] Hemachander Subramanian and Robert A Gatenby. “Evolutionary advantage of directional symmetry breaking in self-replicating polymers”. In: Journal of theoretical biology 446 (2018), pp. 128–136.

[73] Hemachander Subramanian. “Joint optimization of replicative rate and information storage set the letter size of primordial genetic alphabet”. In: BioSystems 251 (2025), p. 105442.

[74] Guillaume Guilbaud et al. “Determination of human DNA replication origin position and efficiency reveals principles of initiation zone organisation”. In: Nucleic Acids Research 50.13 (2022), pp. 7436–7450.

[75] Sjoerd Wanrooij et al. “In vivo mutagenesis reveals that OriL is essential for mitochondrial DNA replication”. In: EMBO reports 13.12 (2012), pp. 1130–1137.

[76] Marie Touchon and Eduardo PC Rocha. “Causes of insertion sequences abundance in prokaryotic genomes”. In: Molecular biology and evolution 24.4 (2007), pp. 969–981.

[77] Sara Vieira-Silva and Eduardo PC Rocha. “The systemic imprint of growth and its uses in ecological (meta) genomics”. In: PLoS genetics 6.1 (2010), e1000808.

[78] Ren Zhang and Chun-Ting Zhang. “Identification of replication origins in archaeal genomes based on the Zcurve method”. In: Archaea 1.5 (2004), p. 335.

[79] Natalia V Sernova and Mikhail S Gelfand. “Identification of replication origins in prokaryotic genomes”. In: Briefings in Bioinformatics 9.5 (2008), pp. 376–391.

[80] AC Frank and JR Lobry. “Oriloc: prediction of replication boundaries in unannotated bacterial chromosomes”. In: Bioinformatics 16.6 (2000), pp. 560–561.

[81] Ren Zhang and Chun-Ting Zhang. “Z curves, an intutive tool for visualizing and analyzing the DNA sequences”. In: Journal of Biomolecular Structure and Dynamics 11.4 (1994), pp. 767–782.

[82] Feng Gao and Chun-Ting Zhang. “Ori-Finder: a webbased system for finding oriC s in unannotated bacterial genomes”. In: BMC bioinformatics 9 (2008), pp. 1–6.

[83] T David Matthews and Stanley Maloy. “Fitness effects of replichore imbalance in Salmonella enterica”. In: Journal of bacteriology 192.22 (2010), pp. 6086–6088.

[84] Aaron E Darling, István Miklós, and Mark A Ragan. “Dynamics of genome rearrangement in bacterial populations”. In: PLoS genetics 4.7 (2008), e1000128.

[85] Magnus G Jespersen et al. “Insertion sequence elements and unique symmetrical genomic regions mediate chromosomal inversions in Streptococcus pyogenes”. In: Nucleic Acids Research 52.21 (2024), pp. 13128–13137.

[86] Chih-Horng Kuo and Howard Ochman. “Deletional bias across the three domains of life”. In: Genome biology and evolution 1 (2009), pp. 145–152.

[87] Martin S Taylor, Chris P Ponting, and Richard R Copley. “Occurrence and consequences of coding sequence insertions and deletions in mammalian genomes”. In: Genome research 14.4 (2004), pp. 555–566.

[88] Shiheng Tao et al. “Patterns of insertion and deletion in mammalian genomes”. In: Current Genomics 8.6 (2007), pp. 370–378.

[89] Jan O Andersson and Siv GE Andersson. “Pseudogenes, junk DNA, and the dynamics of Rickettsia genomes”. In: Molecular biology and evolution 18.5 (2001), pp. 829–839.

[90] T Ryan Gregory. “Insertion–deletion biases and the evolution of genome size”. In: Gene 324 (2004), pp. 15–34.

[91] Erwin Chargaff. “Chemical specificity of nucleic acids and mechanism of their enzymatic degradation”. In: Experientia 6.6 (1950), pp. 201–209.

[92] Erwin Chargaff, Rakoma Lipshitz, and Charlotte Green. “Composition of the desoxypentose nucleic acids of four genera of sea-urchin”. In: Journal of Biological Chemistry 195.1 (1952), pp. 155–160.

[93] Noboru Sueoka. “Intrastrand parity rules of DNA base composition and usage biases of synonymous codons”. In: Journal of molecular evolution 40 (1995), pp. 318– 325.

[94] David Mitchell and Robert Bridge. “A test of Chargaff’s second rule”. In: Biochemical and biophysical research communications 340.1 (2006), pp. 90–94.

[95] Pierre-François Baisnée, Steve Hampson, and Pierre Baldi. “Why are complementary DNA strands symmetric?” In: Bioinformatics 18.8 (2002), pp. 1021–1033.

[96] Donald R Forsdyke. “Neutralism versus selectionism: Chargaff’s second parity rule, revisited”. In: Genetica 149.2 (2021), pp. 81–88.

[97] Donald R Forsdyke. “Relative roles of primary sequence and (G+ C)% in determining the hierarchy of frequencies of complementary trinucleotide pairs in DNAs of different species”. In: Journal of molecular evolution 41 (1995), pp. 573–581.

[98] Shang-Hong Zhang and Ya-Zhi Huang. “Limited contribution of stem-loop potential to symmetry of singlestranded genomic DNA”. In: Bioinformatics 26.4 (2010), pp. 478–485.

[99] JR Lobry. “Properties of a general model of DNA evolution under no-strand-bias conditions”. In: Journal of molecular evolution 40 (1995), pp. 326–330.

[100] Guenter Albrecht-Buehler. “Asymptotically increasing compliance of genomes with Chargaff’s second parity rules through inversions and inverted transpositions”. In: Proceedings of the National Academy of Sciences 103.47 (2006), pp. 17828–17833.

[101] Andrew Hart, Servet Martínez, and Felipe Olmos. “A Gibbs approach to Chargaff’s second parity rule”. In: Journal of Statistical Physics 146 (2012), pp. 408–422.

[102] Piero Fariselli et al. “DNA sequence symmetries from randomness: the origin of the Chargaff’s second parity rule”. In: Briefings in bioinformatics 22.2 (2021), pp. 2172–2181.

[103] Bakhyt T Matkarimov and Murat K Saparbaev. “Chargaff’s second parity rule lies at the origin of additive genetic interactions in quantitative traits to make omnigenic selection possible”. In: PeerJ 11 (2023), e16671.

[104] Patrick Pflughaupt and Aleksandr B Sahakyan. “Generalised interrelations among mutation rates drive the genomic compliance of Chargaff’s second parity rule”. In: Nucleic Acids Research 51.14 (2023), pp. 7409–7423.

[105] Jean-Pierre Vartanian, Michel Henry, and Simon Wain-Hobson. “Hypermutagenic PCR involving all four transitions and a sizeable proportion of transversions”. In: Nucleic acids research 24.14 (1996), pp. 2627–2631.

[106] AC Frank and JR Lobry. “Asymmetric substitution patterns: a review of possible underlying mutational or selective mechanisms”. In: Gene 238.1 (1999), pp. 65–77.

[107] Boris Bartholdy et al. “Allele-specific analysis of DNA replication origins in mammalian cells”. In: Nature communications 6.1 (2015), p. 7051.

[108] Christoforos Nikolaou and Yannis Almirantis. “Deviations from Chargaff’s second parity rule in organellar DNA: Insights into the evolution of organellar genomes”. In: Gene 381 (2006), pp. 34–41.

[109] Yukihiro Kabeya and Shin-ya Miyagishima. “Chloroplast DNA replication is regulated by the redox state independently of chloroplast division in Chlamydomonas reinhardtii”. In: Plant physiology 161.4 (2013), pp. 2102–2112.

[110] Jordan G Okie, Val H Smith, and Mercedes Martin-Cereceda. “Major evolutionary transitions of life, metabolic scaling and the number and size of mitochondria and chloroplasts”. In: Proceedings of the Royal Society B: Biological Sciences 283.1831 (2016), p. 20160611.

[111] Sudhir Kumar et al. “TimeTree 5: an expanded resource for species divergence times”. In: Molecular biology and evolution 39.8 (2022), msac174.

[112] Simon P Blomberg, Theodore Garland Jr, and Anthony R Ives. “Testing for phylogenetic signal in comparative data: behavioral traits are more labile”. In: Evolution 57.4 (2003), pp. 717–745.

[113] José E González-Pastor, José L San Millán, and Felipe Moreno. “The smallest known gene.” In: Nature 369.6478 (1994), pp. 281–281.

[114] John I Glass et al. “Essential genes of a minimal bacterium”. In: Proceedings of the National Academy of Sciences 103.2 (2006), pp. 425–430.

[115] Claire M Fraser et al. “The minimal gene complement of Mycoplasma genitalium”. In: Science 270.5235 (1995), pp. 397–404.

[116] Rosario Gil et al. “Determination of the core of a minimal bacterial gene set”. In: Microbiology and Molecular Biology Reviews 68.3 (2004), pp. 518–537.

[117] Judith K Greenlee, KS Rai, and Alton D Floyd. “Intraspecific variation in nuclear DNA content in Collinsia verna Nutt.(Scrophulariaceae)”. In: Heredity 52.2 (1984), pp. 235–242.

[118] Petr Šmarda and Petr Bureš. “Understanding intraspecific variation in genome size in plants.” In: Preslia 82.1 (2010), pp. 41–61.

[119] Mike D Bennett and Ilia J Leitch. “Nuclear DNA amounts in angiosperms: targets, trends and tomorrow”. In: Annals of botany 107.3 (2011), pp. 467–590.

[120] Ilia J Leitch and Andrew R Leitch. “Genome size diversity and evolution in land plants”. In: Plant genome diversity Volume 2: Physical structure, behaviour and evolution of plant genomes. Springer, 2012, pp. 307–322.

[121] T Ryan Gregory. The evolution of the genome. Elsevier, 2011.

[122] Samuel F Lockwood and John W Bickham. “Genome size in Beaufort Sea coastal assemblages of Arctic ciscoes”. In: Transactions of the American Fisheries Society 121.1 (1992), pp. 13–20.

[123] Dmitri A Petrov. “Mutational equilibrium model of genome size evolution”. In: Theoretical population biology 61.4 (2002), pp. 531–544.

[124] Vinayakumar V Prabhu. “Symmetry observations in long nucleotide sequences.” In: Nucleic acids research 21.12 (1993), p. 2797.

[125] Peyman Fahimi, Chérif F. Matta, and Jordan Okie. “Are Size and Mitochondrial Power of Cells Interdetermined?” In: Journal of Theoretical Biology 572 (July 2023), p. 111565.

[126] Lilas Courtot, Jean-Sébastien Hoffmann, and Valérie Bergoglio. “The protective role of dormant origins in response to replicative stress”. In: International journal of molecular sciences 19.11 (2018), p. 3569.

[127] RA Sclafani and TM Holzen. “Cell cycle regulation of DNA replication”. In: Annu. Rev. Genet. 41.1 (2007), pp. 237–280.

[128] Antoine Aze and Domenico Maiorano. “Recent advances in understanding DNA replication: cell type– specific adaptation of the DNA replication program”. In: F1000Research 7 (2018).

[129] Ichiro Hiratani et al. “Global reorganization of replication domains during embryonic stem cell differentiation”. In: PLoS biology 6.10 (2008), e245.

[130] Tyrone Ryba et al. “Evolutionarily conserved replication timing profiles predict long-range chromatin interactions and distinguish closely related cell types”. In: Genome research 20.6 (2010), pp. 761–770.

[131] David M Gilbert. “Making sense of eukaryotic DNA replication origins”. In: Science 294.5540 (2001), pp. 96– 100.

[132] Nicholas Rhind and David M Gilbert. “DNA replication timing”. In: Cold Spring Harbor perspectives in biology 5.8 (2013), a010132.

[133] Haiqing Fu et al. “Dynamics of replication origin overactivation”. In: Nature Communications 12.1 (2021), p. 3448.

